# The Open Pediatric Cancer Project

**DOI:** 10.1101/2024.07.09.599086

**Authors:** Zhuangzhuang Geng, Eric Wafula, Ryan J. Corbett, Yuanchao Zhang, Run Jin, Krutika S. Gaonkar, Sangeeta Shukla, Komal S. Rathi, Dave Hill, Aditya Lahiri, Daniel P. Miller, Alex Sickler, Kelsey Keith, Christopher Blackden, Antonia Chroni, Miguel A. Brown, Adam A. Kraya, Kaylyn L. Clark, Brian R. Rood, Adam C. Resnick, Nicholas Van Kuren, John M. Maris, Alvin Farrel, Mateusz P. Koptyra, Gerri R. Trooskin, Noel Coleman, Yuankun Zhu, Stephanie Stefankiewicz, Zied Abdullaev, Asif T Chinwalla, Mariarita Santi, Ammar S. Naqvi, Jennifer L. Mason, Carl J. Koschmann, Xiaoyan Huang, Sharon J. Diskin, Kenneth Aldape, Bailey K. Farrow, Weiping Ma, Bo Zhang, Brian M. Ennis, Sarah Tasian, Saksham Phul, Matthew R. Lueder, Chuwei Zhong, Joseph M. Dybas, Pei Wang, Deanne Taylor, Jo Lynne Rokita

**Affiliations:** Center for Data-Driven Discovery in Biomedicine, Children’s Hospital of Philadelphia, Philadelphia, PA, 19104, USA; Division of Neurosurgery, Children’s Hospital of Philadelphia, Philadelphia, PA, 19104, USA; Department of Biomedical and Health Informatics, Children’s Hospital of Philadelphia, Philadelphia, PA, 19104, USA; Center for Cancer and Immunology Research, Children’s National Hospital, Washington, DC, 20010, USA; Center for Data-Driven Discovery in Biomedicine, Children’s Hospital of Philadelphia, Philadelphia, PA, 19104, USA; Division of Neurosurgery, Children’s Hospital of Philadelphia, Philadelphia, PA, 19104, USA; Center for Data-Driven Discovery in Biomedicine, Children’s Hospital of Philadelphia, Philadelphia, PA, 19104, USA; Division of Neurosurgery, Children’s Hospital of Philadelphia, Philadelphia, PA, 19104, USA; Department of Biomedical and Health Informatics, Children’s Hospital of Philadelphia, Philadelphia, PA, 19104, USA; Center for Data-Driven Discovery in Biomedicine, Children’s Hospital of Philadelphia, Philadelphia, PA, 19104, USA; Department of Biomedical and Health Informatics, Children’s Hospital of Philadelphia, Philadelphia, PA, 19104, USA; Center for Cancer and Immunology Research, Children’s National Hospital, Washington, DC, 20010, USA; Center for Cancer and Immunology Research, Children’s National Hospital, Washington, DC, 20010, USA; George Washington University School of Medicine and Health Sciences, Washington, D.C., 20052, USA; Center for Data-Driven Discovery in Biomedicine, Children’s Hospital of Philadelphia, Philadelphia, PA, 19104, USA; Division of Neurosurgery, Children’s Hospital of Philadelphia, Philadelphia, PA, 19104, USA · Funded by Children’s Brain Tumor Network; NIH 3P30 CA016520-44S5, U2C HL138346-03, U24 CA220457-03; NCI/NIH Contract No. 75N91019D00024, Task Order No. 75N91020F00003; Children’s Hospital of Philadelphia Division of Neurosurgery; Division of Oncology, Children’s Hospital of Philadelphia, Philadelphia, PA, 19104, USA; Department of Pediatrics, University of Pennsylvania, Philadelphia, PA, 19104, USA; Department of Biomedical and Health Informatics, Children’s Hospital of Philadelphia, Philadelphia, PA, 19104, USA; Division of Oncology, Children’s Hospital of Philadelphia, Philadelphia, PA, 19104, USA; Center for Childhood Cancer Research, Children’s Hospital of Philadelphia, Philadelphia, PA, 19104, USA · Funded by NCI/NIH Contract No. 75N91019D00024, Task Order No. 75N91020F00003; Laboratory of Pathology, National Cancer Institute, Bethesda, MD, 20892, USA; Department of Pathology and Laboratory Medicine, Children’s Hospital of Philadelphia, Philadelphia, PA, 19104, USA; Department of Pathology and Laboratory Medicine, University of Pennsylvania Perelman School of Medicine, Philadelphia, PA, 19104, USA; Department of Pediatrics, University of Michigan Health, Ann Arbor, MI, 48105, USA; Pediatric Hematology Oncology, Mott Children’s Hospital, Ann Arbor, MI, 48109, USA; Department of Genetics and Genomic Sciences, Icahn School of Medicine at Mount Sinai, New York, NY 10029, USA; Tisch Cancer Institute, Icahn School of Medicine at Mount Sinai, New York, NY, 10029, USA; Center for Data-Driven Discovery in Biomedicine, Children’s Hospital of Philadelphia, Philadelphia, PA, 19104, USA; Division of Neurosurgery, Children’s Hospital of Philadelphia, Philadelphia, PA, 19104, USA; Department of Pathology and Laboratory Medicine, Children’s Hospital of Philadelphia, Philadelphia, PA, 19104, USA; Department of Genetics and Genomic Sciences, Icahn School of Medicine at Mount Sinai, New York, NY 10029, USA; Tisch Cancer Institute, Icahn School of Medicine at Mount Sinai, New York, NY 10029, USA; Department of Biomedical and Health Informatics, Children’s Hospital of Philadelphia, Philadelphia, PA, 19104, USA; Department of Pediatrics, University of Pennsylvania Perelman Medical School, Philadelphia, PA, 19104, USA · Funded by NCI/NIH Contract No. 75N91019D00024, Task Order No. 75N91020F00003; Center for Cancer and Immunology Research, Children’s National Hospital, Washington, DC, 20010, USA; Center for Data-Driven Discovery in Biomedicine, Children’s Hospital of Philadelphia, Philadelphia, PA, 19104, USA; Division of Neurosurgery, Children’s Hospital of Philadelphia, Philadelphia, PA, 19104, USA; Department of Biomedical and Health Informatics, Children’s Hospital of Philadelphia, Philadelphia, PA, 19104, USA · Funded by NCI/NIH Contract No. 75N91019D00024, Task Order No. 75N91020F00003

**Author notes:** Correspondence: Jo Lynne Rokita.

**Keywords:** Pediatric cancer, open science, reproducibility, multi-omics, Docker, OpenPedCan

## Abstract

**Background:** In 2019, the Open Pediatric Brain Tumor Atlas (OpenPBTA) was created as a global, collaborative open-science initiative to genomically characterize 1,074 pediatric brain tumors and 22 patient-derived cell lines. Here, we present an extension of the OpenPBTA called the Open Pediatric Cancer (OpenPedCan) Project, a harmonized open-source multi-omic dataset from 6,112 pediatric cancer patients with 7,096 tumor events across more than 100 histologies. Combined with RNA-Seq from the Genotype-Tissue Expression (GTEx) and The Cancer Genome Atlas (TCGA), OpenPedCan contains nearly 48,000 total biospecimens (24,002 tumor and 23,893 normal specimens).

**Findings:** We utilized Gabriella Miller Kids First (GMKF) workflows to harmonize WGS, WXS, RNA-seq, and Targeted Sequencing datasets to include somatic SNVs, InDels, CNVs, SVs, RNA expression, fusions, and splice variants. We integrated summarized CPTAC whole cell proteomics and phospho-proteomics data, miRNA-Seq data, and have developed a methylation array harmonization workflow to include m-values, beta-vales, and copy number calls. OpenPedCan contains reproducible, dockerized workflows in GitHub, CAVATICA, and Amazon Web Services (AWS) to deliver harmonized and processed data from over 60 scalable modules which can be leveraged both locally and on AWS. The processed data are released in a versioned manner and accessible through CAVATICA or AWS S3 download (from GitHub), and queryable through PedcBioPortal and the NCI’s pediatric Molecular Targets Platform. Notably, we have expanded PBTA molecular subtyping to include methylation information to align with the WHO 2021 Central Nervous System Tumor classifications, allowing us to create research-grade integrated diagnoses for these tumors.

**Conclusions:** OpenPedCan data and its reproducible analysis module framework are openly available and can be utilized and/or adapted by researchers to accelerate discovery, validation, and clinical translation.

## Data Description

The Open Pediatric Cancer (OpenPedCan) project is an iterative open analysis effort in which we harmonize pediatric cancer data from multiple sources, perform downstream cancer analyses on these data, and provide them through Amazon S3, CAVATICA, PedcBioPortal, and v2.1 of NCI’s <u>Pediatric Molecular Targets Platform (MTP)</u>. We harmonized, aggregated, and analyzed data from multiple pediatric and adult data sources, building upon the work of the OpenPBTA (**Figure 1**). All RNA-seq and DNA-seq data from OpenPBTA were updated from GENCODE v27 to GENCODE v39 as part of the OpenPedCan project. Further, all data within OpenPedCan is harmonized with GENCODE v39 annotations. Biospecimen-level metadata and clinical data are contained in **Supplemental Table 1**.

**Figure 1:**
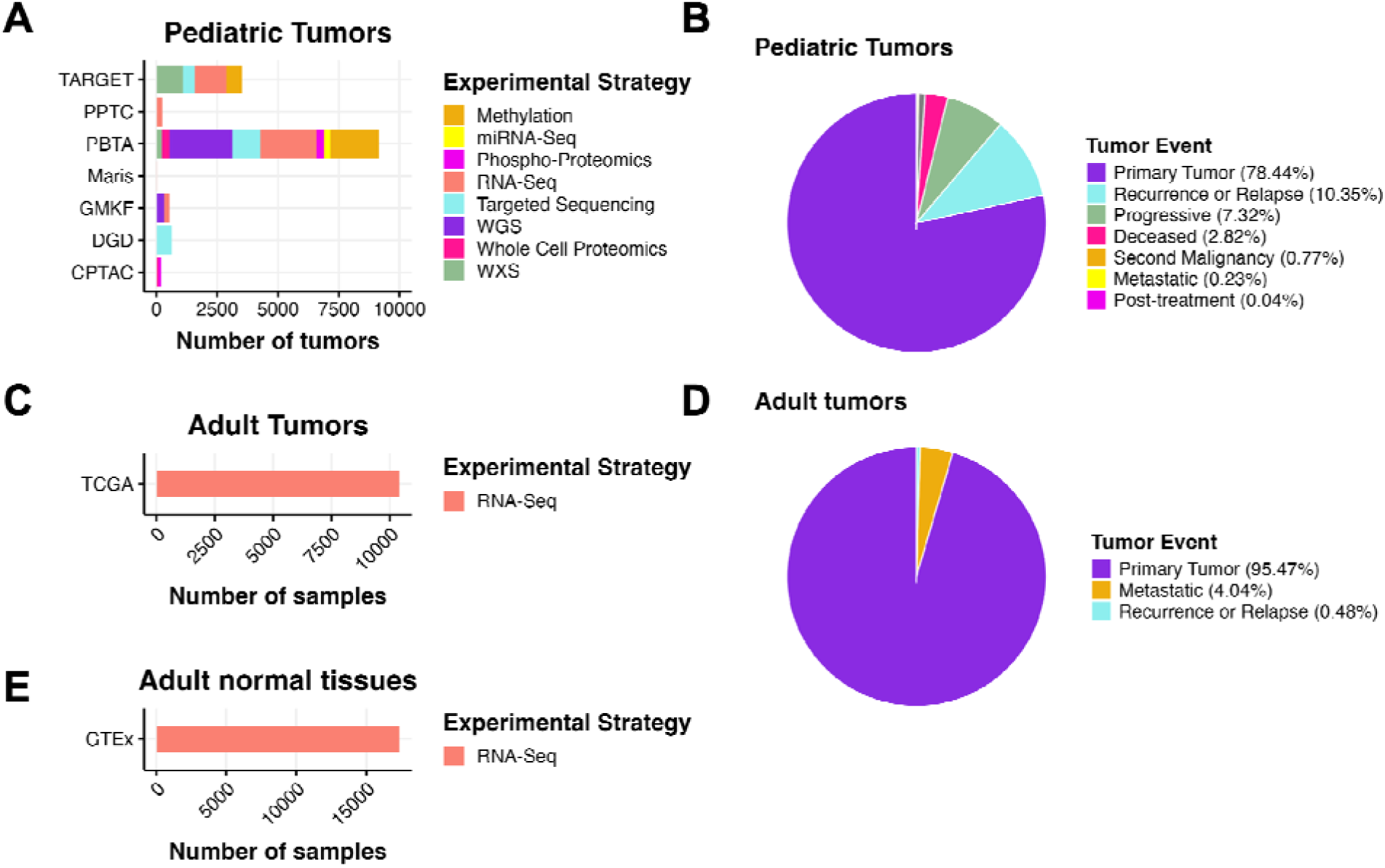
OpenPedCan Data. A, OpenPedCan contains multi-omic data from seven cohorts of pediatric tumors (A-B) with counts by tumor event, RNA-Seq from adult tumors from The Cance Genome Atlas (TCGA) Program (C-D) and RNA-Seq from normal adult tissues from the Genotype-Tissue Expression (GTeX) project (E) with counts by specimen. (Abbreviations: TARGET = Therapeutically Applicable Research to Generate Effective Treatments, PPTC = Pediatric Preclinical Testing Consortium, PBTA = Pediatric Brain Tumor Atlas, Maris = Neuroblastoma cell lines from the Maris Laboratory at CHOP, GMKF = Gabriella Miller Kids First, DGD = Division of Genomic Diagnostics at CHOP, CPTAC = Clinical Proteomic Tumor Analysis Consortium)

OpenPedCan currently include the following datasets, described more fully below:

- OpenPBTA
- TARGET
- Kids First Neuroblastoma (X01)
- Kids First PBTA (X01)
- Chordoma Foundation
- PPTC
- Maris
- MI-ONCOSEQ Study
- DGD
- GTEx
- TCGA
- CPTAC PBTA
- CPTAC GBM
- HOPE proteomics

### Open Pediatric Brain Tumor Atlas (OpenPBTA)

In September of 2018, the <u>Children’s Brain Tumor Network (CBTN)</u> released the <u>Pediatric Brain</u> <u>Tumor Atlas (PBTA)</u>, a genomic dataset (whole genome sequencing, whole exome sequencing, RNA sequencing, proteomic, and clinical data) for nearly 1,000 tumors, available from the <u>Gabriella Miller Kids First Portal</u>. In September of 2019, the Open Pediatric Brain Tumor Atlas (OpenPBTA) Project was launched. OpenPBTA was a global open science initiative to comprehensively define the molecular landscape of tumors of 943 patients from the CBTN and the PNOC003 DIPG clinical trial from the <u>Pediatric Pacific Neuro-oncology Consortium</u> through real-time, collaborative analyses and collaborative manuscript writing on GitHub [1]. Additional PBTA data has been, and will be continually added to, OpenPedCan.

### Therapeutically Applicable Research to Generate Effective Treatments <u>(TARGET)</u>

The Therapeutically Applicable Research to Generate Effective Treatments (TARGET) Initiative is an NCI-funded collection of disease-specific projects that seeks to identify the genomic changes of pediatric cancers. The overall goal is to collect genomic data to accelerate the development of more effective therapies. OpenPedCan analyses include newly harmonized, open-access data associated with the seven diseases present in the TARGET dataset: Acute Lymphoblastic Leukemia (ALL), Acute Myeloid Leukemia (AML), Clear cell sarcoma of the kidney, Neuroblastoma, Osteosarcoma, Rhabdoid tumor, and Wilm’s Tumor.

### Gabriella Miller Kids First <u>(Neuroblastoma)</u> and <u>PBTA</u>

The Gabriella Miller Kids First Pediatric Research Program (Kids First) is a large-scale effort to accelerate research and gene discovery in pediatric cancers and structural birth defects. The program includes whole genome sequencing (WGS) from patients with pediatric cancers and structural birth defects and their families. OpenPedCan analyses include Neuroblastoma and PBTA data from the Kids First projects.

### Chordoma Foundation

The Chordoma Foundation seeks to advance research and improve healthcare for patients diagnosed with chordoma and has shared patient and model sequencing data with the CBTN.

### Pediatric Preclinical Testing Consortium <u>(PPTC)</u>

The National Cancer Institute’s (NCI) former PPTC, now the <u>Pediatric Preclinical in Vivo Testing</u> <u>(PIVOT) Program</u>, molecularly and pharmacologically characterizes cell-derived and patient-derived xenograft (PDX) models. OpenPedCan includes re-harmonized RNA-Seq data for 244 models from the initial PPTC study [2]. A subset of PPTC includes neuroblastoma models; the `Maris` cohort includes re-harmonized RNA-Seq data for 39 neuroblastoma cell lines [3], some of which have corresponding PDX models within the PPTC.

### MI-ONCOSEQ Study [4]

These clinical sequencing data from the University of Michigan were donated to CBTN and added to the PBTA cohort.

### Division of Genomic Diagnostics at Children’s Hospital of Philadelphia <u>(DGD)</u>

CHOP’s Division of Genomic Diagnostics has partnered with CCDI to add somatic panel sequencing data to OpenPedCan and the Molecular Targets Platform.

### The Genotype-Tissue Expression Project <u>(GTEx)</u>

The GTEx project is an ongoing effort to build a comprehensive public data resource and tissue bank to study tissue-specific gene expression, regulation and their relationship with genetic variants. Samples were collected from 54 non-diseased tissue sites across nearly 1000 individuals, primarily for molecular assays including WGS, WXS, and RNA-Seq. OpenPedCan project includes 17,382 GTEx RNA-Seq samples from GTEx v8 release, which span across 31 GTEx groups in the v12 release.

### The Cancer Genome Atlas Program <u>(TCGA)</u>

TCGA is a landmark cancer genomics program that molecularly characterized over 20,000 primary cancer and matched normal samples spanning 33 cancer types. It is a joint effort between NCI and the National Human Genome Research Institute. OpenPedCan project includes open-access 10,414 RNA-Seq for 716 normal and 9,698 TCGA tumor samples from <u>33</u> <u>cancer types</u>.

### Clinical Proteomic Tumor Analysis Consortium (CPTAC) PBTA proteomics study

The CPTAC pediatric pan-brain tumor study [5] contains 218 tumors profiled by proteogenomics and are included in OPC.

### CPTAC adult GBM proteomics study

This CPTAC adult GBM study [6] contains 99 tumors profiled by proteogenomics and are included in OPC.

### Project HOPE proteomics study

Project HOPE is an adolescent and young adult high-grade glioma study (in preparation for publication) that contains 90 tumors profiled by proteogenomics and are included in OPC.

OpenPedCan represents a substantial expansion since the OpenPBTA, both in cohort size and in data modality integration. By incorporating methylation, proteomics, splicing, and reference datasets, and enabling reproducible analyses across more than 48,000 biospecimens, OpenPedCan delivers a uniquely scalable and reusable resource for pediatric cancer research.

## Context

Creation of this dataset had multiple motivations. First, we sought to harmonize, summarize, and contextualize pediatric cancer genomics data among normal tissues (GTEx) and adult cancer tissues (TCGA) to enable the creation of the National Cancer Institute’s Molecular Targets Platform (MTP) at https://moleculartargets.ccdi.cancer.gov/. The inclusion of harmonized GTEx and adult TCGA data specifically allows for the identification of genes and/or transcripts expressed in a tumor-specific and/or pediatric tumor-specific manner. Next, we created this resource for broad community use to promote rapid reuse and accelerate the discovery of additional mechanisms contributing to the pathogenesis of pediatric cancers and/or to identify novel candidate therapeutic targets for pediatric cancer.

Similar to OpenPBTA, OpenPedCan operates on a pull request model to accept contributions. We set up continuous integration software via GitHub Actions to confirm the reproducibility of analyses within the project’s Docker container. We maintained a data release folder on Amazon S3, downloadable directly from S3 or our open-access CAVATICA project, with merged files for each analysis. As we produced new results, identified data issues, or added additional data, we created new data releases in a versioned manner. The project maintainers have included engineers and scientists from the <u>Children’s Hospital of Philadelphia</u> and <u>Children’s National</u> <u>Hospital</u>.

## Methods

An overview of the OpenPedCan methods is depicted in **Figure 2**. Briefly, most primary data harmonization analysis workflows were performed with Kids First pipelines written in Common Workflow Language (CWL) using CAVATICA (detailed below). Alignment and expression quantification for GTEx and TCGA RNA-Seq was performed by the respective consortium.

**Figure 2:**
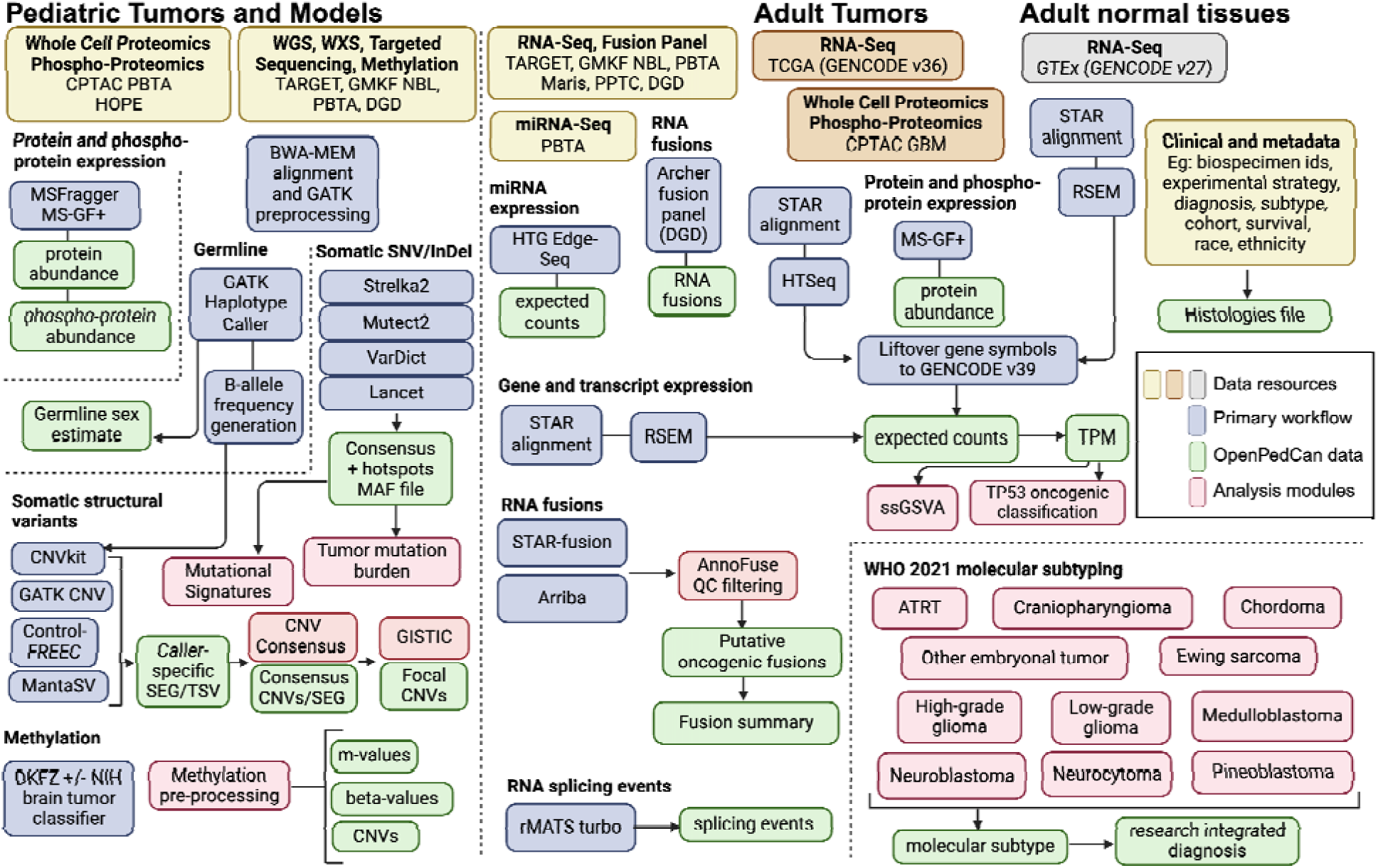
OpenPedCan Analysis Workflow. Depicted are the datasets (yellow, orange, and grey) contained within OpenPedCan. These datasets are made available in a harmonized manner through primary analysis workflows (blue) for DNA, RNA, and/or proteogenomics data. Files derived from the primary analysis workflows (green) are released within OpenPedCan. Additional analysis modules developed within OpenPedCan (red) also generate results files (green) which are released within OpenPedCan.

Custom python, R, and/or bash scripts were then created in OpenPedCan using the primary harmonized output files.

## Sample Details

A list of all biospecimens and associated metadata can be found in **Supplemental Table 1**.

### Nucleic acids extraction and library preparation (PBTA X01 and miRNA-Seq)

For detailed methods about the OpenPBTA cohort, please refer to the manuscript [1]. For the PBTA X01 cohort, libraries were prepped using the Illumina TruSeq Strand-Specific Protocol to pull out poly-adenylated transcripts.

#### cDNA Library Construction

Total RNA was quantified using the Quant-iT RiboGreen RNA Assay Kit and normalized to 5ng/ul. Following plating, 2 uL of ERCC controls (using a 1:1000 dilution) were spiked into each sample. An aliquot of 325 ng for each sample was transferred into library preparation. The resultant 400bp cDNA went through dual-indexed library preparation: ‘A’ base addition, adapter ligation using P7 adapters, and PCR enrichment using P5 adapters. After enrichment, the libraries were quantified using Quant-iT PicoGreen (1:200 dilution). Samples were normalized to 5 ng/uL. The sample set was pooled and quantified using the KAPA Library Quantification Kit for Illumina Sequencing Platforms.

#### miRNA Extraction and Library Preparation

Total RNA for CBTN samples was extracted as described in OpenPBTA [1] and prepared according to the HTG Edge Seq protocol for the extracted RNA miRNA Whole transcriptome assay (WTA). 15ng of RNA were mixed in 25ul of lysis buffer, which were then loaded onto a 96-well plate. Human Fetal Brain Total RNA (Takara Bio USA, #636526) and Human Brain Total RNA (Ambion, Inc., Austin, TX, USA) were used as controls. The plate was loaded into the HTG EdgeSeq processor along with the miRNA WTA assay reagent pack. Samples were processed for 18-20 hours, then were barcoded and amplified using a unique forward and reverse primer combination. PCR settings used for barcoding and amplification were 95C for 4 min, 16 cycles of (95C for 15 sec, 56C for 45 sec, 68C for 45 sec), and 68C for 10 min. Barcoded and amplified samples were cleaned using AMPure magnetic beads (Ampure XP,Cat# A63881). Libraries were quantified using the KAPA Biosystem assay qPCR kit (Kapa Biosystems Cat#KK4824) and CT values were used to determine the pM concentration of each library.

### Data generation

#### PBTA X01 Illumina Sequencing

Pooled libraries were normalized to 2nM and denatured using 0.1 N NaOH prior to sequencing. Flowcell cluster amplification and sequencing were performed according to the manufacturer’s protocols using the NovaSeq 6000. Each run was a 151bp paired-end with an eight-base index barcode read. Data was analyzed using the Broad Picard Pipeline which includes de-multiplexing and data aggregation.

#### PBTA miRNA Sequencing

Libraries were pooled, denatured, and loaded onto sequencing cartridge. Libraries were sequenced using an Illumina Nextseq 500 per manufacturer guidelines. FASTQ files were generated from raw sequencing data using Illumina BaseSpace and analyzed with the HTG EdgeSeq Parser software v5.4.0.7543 to generate an excel file containing quantification of 2083 miRNAs per sample. Any sample that did not pass the quality control set by the HTG REVEAL software version 2.0.1 (Tuscon, AR, USA) was excluded from the analysis.

### Primary Workflows through Kids First

#### DNA WGS Alignment and SNP Calling

Please refer to the OpenPBTA manuscript for details on DNA WGS Alignment, prediction of participants’ genetic sex, and SNP calling for B-allele Frequency (BAF) generation. [1].

#### Somatic Mutation and INDEL Calling

For matched tumor/normal samples, we used the same mutation calling methods as described in OpenPBTA manuscript for details [1]. For tumor only samples, we ran Mutect2 from GATK v4.2.2.0 using the following <u>workflow</u>.

##### VCF annotation and MAF creation

Somatic variants were annotated by the Ensembl Variant Effect Predictor (VEP v105) [7]. From tumor only variant calls, we removed variants with `alt_depth == 0` or `t_depth < 4`.

##### Consensus SNV Calling (tumor/normal only)

We adopted the consensus SNV calling method described in OpenPBTA manuscript with adjustment [1]. For SNV calling, we combined four consensus SNV calling algorithms: Strelka2[8], Mutect2[9], Lancet[10], and VarDict[11].

Strelka2 outputs multi-nucleotide polymorphisms (MNPs) as consecutive single-nucleotide polymorphisms. In order to preserve MNPs, we gather MNP calls from the other caller inputs, and search for evidence supporting these consecutive SNP calls as MNP candidates. Once found, the Strelka2 SNP calls supporting a MNP are converted to a single MNP call. This is done to preserve the predicted gene model as accurately as possible in our consensus calls. Consensus SNV from all four callers were collected and by default, calls that were detected in at least two calling algorithms or marked with “HotSpotAllele” were retained.

For all SNVs, potential non-hotspot germline variants were removed if they had a normal depth <= 7 and gnomAD allele frequency > 0.001. Final results were saved in MAF format.

### Somatic Copy Number Variant (CNV) Calling

We called copy number variants for tumor/normal samples using Control-FREEC [12,13] and CNVkit [14] as described in the OpenPBTA manuscript [1]. We used GATK [15] to call CNVs for matched tumor/normal WGS samples when there were at least 30 male and 30 female normals from the same sequencing platform available for panel of normal creation. For tumor only samples, we used Control-FREEC with the following modifications. Instead of the b-allele frequency germline input file, we used the `dbSNP_v153_ucsc-compatible.converted.vt.decomp.norm.common_snps.vcf.gz` <u>dbSNP common snps</u> <u>file</u> and to avoid hard-to-call regions, utilized the `hg38_canonical_150.mappability` <u>mappability file</u>. Both are also linked in the public <u>Kids First references CAVATICA project</u>. The Control-FREEC tumor only workflow can be found <u>here</u>.

### Somatic Structural Variant Calling (WGS samples only)

We called structural variants (SVs) using Manta [16], restricting analysis to the same regions utilized by Strelka2. We annotated SVs using AnnotSV [17].

### Gene Expression

The `tumor-normal-differential-expression` module performs differential expression analyses for all sets of Disease (`cancer_group`) and Dataset (`cohort`) across all genes found in the `gene-expression-rsem-tpm-collapsed.rds` table. The purpose of this analysis is to highlight the correlation and understand the variability in gene expression in different cancer conditions across different histological tissues. For OpenPedCan v12 data release, this module performs expression analysis over 102 cancer groups across 52 histological tissues for all 54,346 genes found in the dataset. This analysis was performed on the Children’s Hospital of Philadelphia HPC and was configured to use 96G of RAM per CPU, with one task (one iteration of expression analysis for each set of tissue and cancer group) per CPU (total 102×52=5304 CPUs) using the <u>R/DESeq2</u> package. Please refer to script `run-tumor-normal-differential-expression.sh` in the module for additional details on Slurm processing configuration. The same analysis can also be performed on CAVATICA, but requires further optimization. The module describes the steps for CAVATICA set up, and scripts to publish an application on the portal. The required data files are also available publicly on CAVATICA under the <u>Open Pediatric Cancer (OpenPedCan) Open Access</u>. Refer to the module for detailed description and scripts.

#### Abundance Estimation

Among the data sources used for OpenPedCan, GTEx and TCGA used GENCODE v27 and v36, respectively. Therefore, the gene symbols had to be harmonized to GENCODE v39 for compatibility with the rest of the dataset. The liftover process was done via a <u>custom script</u>. The script first constructs an object detailing the gene symbol changes from the <u>HGNC symbol</u> <u>database</u>. Using the symbol-change object, the script updates any columns containing gene symbols. This liftover process was used on GTEx RNA-Seq, TCGA RNA-Seq, DGD fusions, and DNA hotspot files.

Additionally, the gene expression matrices had some instances where multiple Ensembl gene identifiers mapped to the same gene symbol. This was dealt with by filtering the expression matrix to only genes with [FPKM/TPM] > 0 and then selecting the instance of the gene symbol with the maximum mean [FPKM/TPM/Expected_count] value across samples. This enabled many downstream modules that require RNA-seq data have gene symbols as unique gene identifiers. Refer to <u>collapse-rnaseq</u> module for scripts and details.

#### Gene fusion detection from RNA-Seq

Gene fusions were called using Arriba [18] and STAR-Fusion [19] as previously reported in OpenPBTA [1]. We updated the `annoFuseData` <u>R package</u> to liftover gene symbols to be concordant with VEP v105. Fusions are now filtered with annoFuse [20] upstream and released in `fusion-annoFuse.tsv.gz`.

#### Gene fusion detection from fusion panels (DGD only)

Clinical RNA fusion calls from the CHOP DGD fusion panel are included in the data release in the fusion-dgd.tsv.gz file.

### Splicing quantification

To detect alternative splicing events, we utilized rMATS turbo (v. 4.1.0) with Ensembl/GENCODE v39 GFF annotations using the <u>Kids First RNA-Seq workflow</u>. We used `--variable-read-length` and `-t paired` options and applied an additional filter to include only splicing events with total junction read counts greater than 10. The OpenPedCan data release file `splice-events-rmats.tsv.gz` contains predicted single exon (SE), alternative 5’ splice site (A5SS), alternative 3’ splice site (A3SS), and retained intron (RI) events. These are made available for the community, but were not yet used in OpenPedCan analysis modules.

### Proteomics data integration

#### CPTAC PBTA, CPTAC GBM, and HOPE proteogenomics

The following methods are the general proteomics approaches used for the CPTAC PBTA [5], CPTAC GBM [6], and HOPE (pre-publication, correspondence with Dr. Pei Wang) studies. For specific descriptions of sample preparation, mass spectrometry instrumentation and approaches, and data generation, processing, or analysis please refer to the relevant publications.

##### TMT-11 Labeling and Phosphopeptide Enrichment

Proteome and phosphoproteome analysis of brain cancer samples in the CPTAC PBTA (pediatric), CPTAC GBM (adult), and HOPE (adolescent and young adult, AYA) cohort studies were structured as TMT11-plex experiments. Tumor samples were digested with LysC and trypsin. Digested peptides were labeled with TMT11-plex reagent and prepared for phosphopeptide enrichment. For each dataset, a common reference sample was compiled from representative samples within the cohort. Phosphopeptides were enriched using Immobilized Metal Affinity Chromatography (IMAC) with Fe3+-NTA-agarose bead kits.

##### Liquid Chromatography with Tandem Mass Spectrometry (LC-MS/MS) Analysis

To reduce sample complexity, peptide samples were separated by high pH reversed phase HPLC fractionation. For CPTAC PBTA a total of 96 fractions were consolidated into 12 final fractions for LC-MS/MS analysis. For CPTAC GBM and HOPE cohorts a total of 96 fractions were consolidated into 24 fractions. For CPTAC PBTA, global proteome mass spectrometry analyses were performed on an Orbitrap Fusion Tribrid Mass Spectrometer and phosphoproteome analyses were performed on an Orbitrap Fusion Lumos Tribrid Mass Spectrometer. For CPTAC GBM and HOPE studies, mass spectrometry analysis was performed using an Orbitrap Fusion Lumos Mass Spectrometer.

##### Protein Identification

The CPTAC PBTA spectra data were analyzed with MSFragger version 20190628 [21] searching against a CPTAC harmonized RefSeq-based sequence database containing 41,457 proteins mapped to the human reference genome (GRCh38/hg38) obtained via the UCSC Table Browser on June 29, 2018, with the addition of 13 proteins encoded in the human mitochondrial genome, 264 common laboratory contaminant proteins, and an equal number of decoy sequences. The CPTAC GBM and HOPE spectra data were analyzed with MS-GF+ v9881 [22,23,24] searching against the RefSeq human protein sequence database downloaded on June 29, 2018 (hg38; 41,734 proteins), combined with 264 contaminants, and a decoy database composed of the forward and reversed protein sequences.

##### Protein Quantification and Data Analysis

Relative protein (gene) abundance was calculated as the ratio of sample abundance to reference abundance using the summed reporter ion intensities from peptides mapped to the respective gene. For phosphoproteomic datasets, data were not summarized by protein but left at the phosphopeptide level. Global normalization was performed on the gene-level abundance matrix (log2 ratio) for global proteomic and on the site-level abundance matrix (log2 ratio) for phosphoproteomic data. The median, log2 relative protein or peptide abundance for each sample was calculated and used to normalize each sample to achieve a common median of 0. To identify TMT outliers, inter-TMT t-tests were performed for each individual protein or phosphopeptide. Batch effects were checked using the log2 relative protein or phosphopeptide abundance and corrected using the Combat algorithm [25]. Imputation was performed after batch effect correction for proteins or phosphopeptides with a missing rate < 50%. For the phosphopeptide datasets, 440 markers associated with cold-regulated ischemia genes were filtered and removed.

### Creation of OpenPedCan Analysis modules

A list of all modules, repository links, one line description, input, and output files can be found in **<u>Supplemental Table 2</u>**.

### Methylation Analysis

#### Methylation array preprocessing

We preprocessed raw Illumina 450K and EPIC 850K Infinium Human Methylation Bead Array intensities using the array preprocessing methods implemented in the `minfi Bioconductor package` [26]. We utilized either `preprocessFunnorm` when an array dataset had both tumor and normal samples or multiple OpenPedCan-defined `cancer_groups` and `preprocessQuantile` when an array dataset had only tumor samples from a single OpenPedCan-defined `cancer_group` to estimate usable methylation measurements (`beta-values` and `m-values`) and copy number (`cn-values`). Some Illumina Infinium array probes targeting CpG loci contain single-nucleotide polymorphisms (SNPs) near or within the probe [27], which could affect DNA methylation measurements [28]. As the minfi preprocessing workflow recommends, we dropped probes containing common SNPs in dbSNP (minor allele frequency > 1%) at the CpG interrogation or the single nucleotide extensions.

Details of methylation array preprocessing are available in the OpenPedCan methylation-preprocessing module.

#### Methylation classification of brain tumor molecular subtypes

The Clinical Methylation Unit Laboratory of Pathology at the National Cancer Institute Center for Cancer Research ran the <u>DKFZ brain classifier version 12.6</u>, a comprehensive DNA methylation-based classification of CNS tumors across all entities and age groups [29] and/or the NIH Bethesda Brain tumor classifier v2.0 (NIH_v2) and the combo reporter pipeline v2.0 on docker container trust1/bethesda:latest. Unprocessed IDAT-files from the <u>Children’s Brain</u> <u>Tumor Network (CBTN)</u> `Infinium Human Methylation EPIC (850k) BeadChip arrays` were used as input and the following information was compiled into the `histologies.tsv` file: `dkfz_v12_methylation_subclass` (predicted methylation subtype), `dkfz_v12_methylation_subclass_score` (classification score), `dkfz_v12_methylation_mgmt_status` (*MGMT* methylation status), `dkfz_v12_methylation_mgmt_estimated` (estimated *MGMT* methylation fraction), `NIH_v2_methylation_Superfamily`, `NIH_v2_methylation_Superfamily_mean_score`, `NIH_v2_methylation_Superfamily_Consistency_score`, `NIH_v2_methylation_Class`, `NIH_v2_methylation_Class_mean_score`, `NIH_v2_methylation_Class_consistency_score`, `NIH_v2_methylation_Superfamily_match`, and `NIH_v2_methylation_Class_match`.

#### Gene Set Variation Analysis (gene-set-enrichment-analysis analysis module)

We performed Gene Set Variation Analysis (GSVA) for the Hallmark gene sets from MSigDB [30] on log2-transformed, gene-collapsed RSEM TPM expression values from RNA-Seq using the GSVA package from Bioconductor [31]. GSVA was performed separately by RNA library type to avoid batch effects.

#### Fusion prioritization (fusion_filtering analysis module)

The `fusion_filtering` module filters artifacts and annotates fusion calls, with prioritization for oncogenic fusions, for the fusion calls from STAR-Fusion and Arriba. After artifact filtering, fusions were prioritized and annotated as “putative oncogenic fusions” when at least one gene was a known kinase, oncogene, tumor suppressor, curated transcription factor, on the COSMIC Cancer Gene Census List, or observed in TCGA. Fusions were retained in this module if they were called by both callers, recurrent or specific to a cancer group, or annotated as a putative oncogenic fusion. Please refer to the module linked above for more detailed documentation and scripts.

#### Consensus CNV Calling (WGS samples only) (copy_number_consensus_call* analysis modules)

We adopted the consensus CNV calling described in OpenPBTA manuscript [1] with minor adjustments. For each caller and sample with WGS performed, we called CNVs based on consensus among Control-FREEC [12,13], CNVkit [14], and GATK [15]. Sample and consensus caller files with more than 2,500 CNVs were removed to de-noise and increase data quality, based on cutoffs used in GISTIC [32]. For each sample, we included the following regions in the final consensus set: 1) regions with reciprocal overlap of 50% or more between at least two of the callers; 2) smaller CNV regions in which more than 90% of regions were covered by another caller. For GATK, if a panel of normal was not able to be created (required 30 male and 30 female with the same sequencing platform), consensus was run for that tumor using Control-FREEC, CNVkit, and MantaSV. We defined copy number as NA for any regions that had a neutral call for the samples included in the consensus file. We merged CNV regions within 10,000 bp of each other with the same direction of gain or loss into single region.

Any CNVs that overlapped 50% or more with immunoglobulin, telomeric, centromeric, segment duplicated regions, or that were shorter than 3000 bp were filtered out. The CNVKit calls for WXS samples were appended to the consensus CNV file.

#### Focal Copy Number Calling (focal-cn-file-preparation analysis module)

Please refer to the OpenPBTA manuscript for details on assignment of copy number status values to CNV segments, cytobands, and genes [1]. We applied criteria to resolve instances of multiple conflicting status calls for the same gene and sample, which are described in detail in the <u>focal-cn-file-preparation</u> module. Briefly, we prioritized 1) non-neutral status calls, 2) calls made from dominant segments with respect to gene overlap, and 3) amplification and deep deletion status calls over gain and loss calls, respectively, when selecting a dominant status call per gene and sample. These methods resolved >99% of duplicated gene-level status calls.

#### Mutational Signatures (mutational-signatures analysis module)

We obtained mutational signature weights (i.e., exposures) from consensus SNVs using the deconstructSigs R package [33]. We estimated weights for single- and double-base substitution (SBS and DBS, respectively) signatures from the Catalogue of Somatic Mutations in Cancer (COSMIC) database versions 2 and 3.3, as well as SBS signatures from Alexandrov et al. 2013 [34]. The following COSMIC SBS signatures were excluded from weight estimation in all tumors: sequencing artifact signatures, 2) signatures associated with environmental exposure, and 3) signatures with an unknown etiology. Additionally, we excluded therapy-associated signatures from mutational signature weight estimation in tumors collected prior to treatment (i.e. “Initial CNS Tumor” or “Primary Tumor”).

#### Tumor Mutation Burden [TMB] (tmb-calculation analysis module)

Recent clinical studies have associated high TMB with improved patient response rates and survival benefit from immune checkpoint inhibitors [35].

The <u>Tumor Mutation Burden (TMB)</u> `tmb-calculation` module was adapted from the `snv-callers` <u>module</u> of the OpenPBTA project [1]. Here, we use mutations in the `snv-consensus-plus-hotspots.maf.tsv.gz` file which is generated using <u>Kids First DRC</u> <u>Consensus Calling Workflow</u> and is included in the OpenPedCan data download. The consensus MAF contains SNVs or MNVs called in at least 2 of the 4 callers (Mutect2, Strelka2, Lancet, and Vardict) plus hotspot mutations if called in 1 of the 4 callers. We calculated TMB for tumor samples sequenced with either WGS or WXS. Briefly, we split the SNV consensus MAF into SNVs and multinucleotide variants (MNVs). We split the MNV subset into SNV calls, merged those back with the SNVs subset, and then removed sample-specific redundant calls. The resulting merged and non-redundant SNV consensus calls were used as input for the TMB calculation. We tallied only nonsynonymous variants with classifications of high/moderate consequence (“Missense_Mutation”, “Frame_Shift_Del”, “In_Frame_Ins”, “Frame_Shift_Ins”, “Splice_Site”, “Nonsense_Mutation”, “In_Frame_Del”, “Nonstop_Mutation”, and “Translation_Start_Site”) for the numerator. All BED files are provided in the data release.

##### All mutation TMB

For WGS samples, we calculated the size of the genome covered as the intersection of Strelka2 and Mutect2’s effectively surveyed areas, regions common to all variant callers, and used this as the denominator. `WGS_all_mutations_TMB = (total # mutations in consensus MAF) / intersection_strelka_mutect_vardict_genome_size` For WXS samples, we used the size of the WXS bed region file as the denominator. `WXS_all_mutations_TMB = (total # mutations in consensus MAF)) / wxs_genome_size`

##### Coding only TMB

We generated coding only TMB from the consensus MAF as well. We calculated the intersection for Strelka2 and Mutect2 surveyed regions using the coding sequence ranges in the GENCODE v39 gtf supplied in the OpenPedCan data download. We removed SNVs outside of these coding sequences prior to implementing the TMB calculation below: `WGS_coding_only_TMB = (total # coding mutations in consensus MAF) / intersection_wgs_strelka_mutect_vardict_CDS_genome_size` For WXS samples, we intersected each WXS bed region file with the GENCODE v39 coding sequence, sum only variants within this region for the numerator, and calculate the size of this region as the denominator. `WXS_coding_only_TMB = (total # coding mutations in consensus MAF) / intersection_wxs_CDS_genome_size` Finally, we include an option (`nonsynfilter_focr`) to use specific nonsynonymous mutation variant classifications recommended from the <u>TMB Harmonization Project</u>.

#### Molecular Subtyping

Here, we build upon the molecular subtyping performed in OpenPBTA [1] to align with WHO 2021 subtypes [36]. Molecular subtypes were generated per tumor event and are listed for each biospecimen in **Supplemental Table 1**, with the number of tumors grouped by broad histology and molecular subtype in **Supplemental Table 3**.

##### High-grade gliomas

High-grade gliomas (HGG) were categorized based on a combination of clinical information, molecular features, and DNA methylation data. H3 K28-altered diffuse midline gliomas (DMG) were classified based on the presence of a p.K28M or p.K28I mutation in *H3F3A*, *HIST1H3B*, *HIST1H3C*, or *HIST2H3C*, or a high-confidence DKFZ methylation score (>=0.8) in the appropriate subclass. Oligodendroglioma, IDH-mutant tumors were classified based on high-confidence “O_IDH” methylation classifications, and oligosarcoma, IDH-mutant tumors were defined as those with high-confidence “OLIGOSARC_IDH” methylation classifications. Pleomorphic xanthoastrocytomas (PXA) were classified using the following criteria: 1) methylation subtype is high-confidence “PXA” or `pathology_free_text_diagnosis` contains “pleomorphic xanthoastrocytoma” or “pxa”, and 2) tumor contains a BRAF V600E mutation and a *CDKN2A* or *CDKN2B* homozygous deletion. Methylation classifications were used in classifying the following subtypes:

1. DHG, H3 G35 (“DHG_G34” and “GBM_G34” classifications)
2. HGG, IDH (“A_IDH_HG” and “GBM_IDH” classifications)
3. HGG, H3 wild type (methylation classification contains “GBM_MES”, “GBM_RTK”, “HGG_”, “HGAP”, “AAP”, or “ped_”)

A new high-grade glioma entity called infant-type hemispheric gliomas (IHGs), characterized by distinct gene fusions enriched in receptor tyrosine kinase (RTK) genes including *ALK*, *NTRK1/2/3*, *ROS1* or *MET*, was identified in 2021 [37]. To identify IHG tumors, first, tumors which were classified as “IHG” by the DKFZ methylation classifier or diagnosed as “infant type hemispheric glioma” from `pathology_free_text_diagnosis` were selected [29]. Then, the corresponding tumor RNA-seq data were utilized to seek the evidence for RTK gene fusion. Based on the specific RTK gene fusion present in the samples, IHGs were further classified as “IHG, ALK-altered”, “IHG, NTRK-altered”, “IHG, ROS1-altered”, or “IHG, MET-altered”. If no fusion was observed, the samples were identified as “IHG, To be classified”.

##### Atypical teratoid rhabdoid tumors

Atypical teratoid rhabdoid tumors (ATRT) tumors were categorized into three subtypes: “ATRT, MYC”, “ATRT, SHH”, and “ATRT, TYR” [38]. In OpenPedCan, the molecular subtyping of ATRT was based solely on the DNA methylation data. Briefly, ATRT samples with a high confidence DKFZ methylation subclass score (>= 0.8) were selected and subtypes were assigned based on the DKFZ methylation subclass [29]. Samples with low confidence DKFZ methylation subclass scores (< 0.8) were identified as “ATRT, To be classified”.

##### Neuroblastoma tumors

Neuroblastoma (NBL) tumors with a pathology diagnosis of neuroblastoma, ganglioneuroblastoma, or ganglioneuroma were subtyped based on their MYCN copy number status as either “NBL, MYCN amplified” or “NBL, MYCN non-amplified”. If `pathology_free_text_diagnosis` was “NBL, MYCN non-amplified” and the genetic data suggested MYCN amplification, the samples were subtyped as “NBL, MYCN amplified”. On the other hand, if `pathology_free_text_diagnosis` was “NBL, MYCN amplified” and the genetic data suggested MYCN non-amplification, the RNA-Seq gene expression level of *MYCN* was used as a prediction indicator. In those cases, samples with *MYCN* gene expression above or below the cutoff (TPM >= 140.83 based on visual inspection of MYCN CNV status) were subtyped as “NBL, MYCN amplified” and “NBL, MYCN non-amplified”, respectively. *MYCN* gene expression was also used to subtype samples without DNA sequencing data. If a sample did not fit none of these situations, it was denoted as “NBL, To be classified”.

##### Craniopharyngiomas

In addition to molecular criteria established in OpenPBTA [1], craniopharyngiomas (CRANIO) are now subtyped using DNA methylation classifiers. Craniopharyngiomas with a high-confidence methylation subclass containing “CPH_PAP” were classified as papillary (CRANIO, PAP), and those with high-confidence methylation subclass containing “CPH_ADM” were classified as adamantinomatous (CRANIO, ADAM), respectively.

##### Ependymomas

Ependymomas (EPN) are subtyped using the following criteria:

1. Any spinal tumor with *MYCN* amplification or with a high-confidence “EPN, SP-MYCN” methylation classification was subtyped as EPN, spinal and MYCN-amplified (SP-MYCN).
2. EPN tumors containing one or more gene fusions of *YAP1::MAMLD1*, *YAP1::MAML2*, or *YAP1::FAM118B*, or else had a high-confidence “EPN, ST YAP1” methylation classification were subtyped as EPN, ST YAP1.
3. EPN tumors containing one or more gene fusions of *ZFTA::RELA* or *ZFTA::MAML2*, or else had a high-confidence “EPN, ST ZFTA” methylation classification were subtyped as EPN, ST ZFTA. This reflects an update to WHO classifications that now characterizes this subtype based on *ZFTA* fusions rather than *RELA* fusions.
4. EPN tumors with 1) chromosome 1q gain and *TKTL1* over-expression, or 2) *EZHIP* over-expression, or 3) posterior fossa anatomical location and a histone H3 K28 mutation in *H3F3A*, *HIST1H3B*, *HIST1H3C*, or *HIST2H3C*, or 4) a high-confidence “EPN, PF A” methylation classification were subtyped as posterior fossa group A ependymomas (EPN, PF A).
5. Tumors with 1) chr 6p or 6q loss and *GPBP1* or *IFT46* over-expression, or 2) a high-confidence “EPN, PF B” methylation classification were subtyped as posterior fossa group B ependymomas (EPN, PF B).
6. EPN tumors with a high-confidence “EPN, MPE” methylation classification were subtyped as myxopapillary ependymomas (EPN, MPE).
7. EPN tumors with a high-confidence “EPN, PF SE” methylation classification were subtyped as posterior fossa subependymomas (EPN, PF SE).
8. EPN tumors with a high-confidence “EPN, SP SE” methylation classification were subtyped as spinal subependymomas (EPN, SP SE).
9. EPN tumors with a high-confidence “EPN, SP” methylation classification were subtyped as spinal ependymomas (EPN, SP).
10. All other EPN tumors were classified as “EPN, To be classified”.

##### Low-grade gliomas

In addition to subtyping methods described in OpenPBTA [1], high-confidence methylation classifications are now used in classifying the following low-grade glioma (LGG) subtypes:

1. LGG, other MAPK-altered (methylation subclass “PA_MID” or “PLNTY”)
2. LGG, FGFR-altered (methylation subclass “PA_INF_FGFR”)
3. LGG, IDH-altered (methylation subclass “A_IDH_LG”)
4. LGG, MYB/MYBL1 fusion (methylation subclass “AG_MYB” or “LGG_MYB”)
5. LGG, MAPK-altered (methylation subclass “LGG, MAPK”)
6. LGG, BRAF- and MAPK-altered (methylation subclass “LGG, BRAF/MAPK”)
7. SEGA, to be classified (methylation subclass “SEGA, To be classified”)

##### Medulloblastomas (MBs)

In addition to our previous work classifying MB tumors into the four major subtypes (WNT, SHH, Group 3, and Group 4) using the transcriptomic MedulloClassifier [39], we integrated high-confidence methylation classification, demographic, and molecular criteria to molecularly subtype SHH tumors into one of four subgroups (alpha, beta, gamma, or delta) (**Figure 3**).

**Figure 3:**
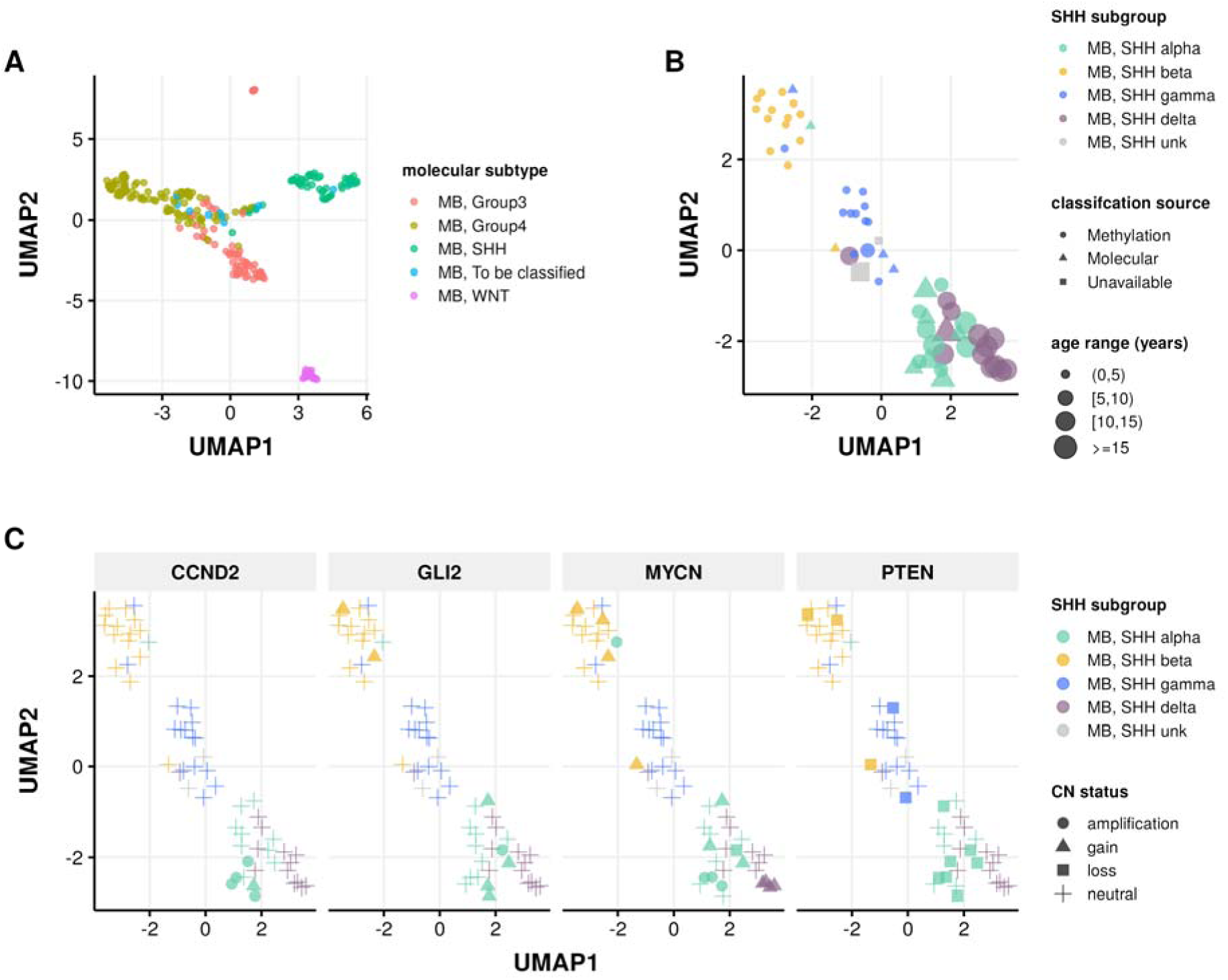
Medulloblastoma Sample Clustering. A, UMAP projection of 271 MB tumors and B, 63 SHH-activated MB tumors using methylation beta values of the 20,000 most variable probes from the Infinium MethylationEPIC array. C, UMAP projection of MB, SHH activated samples indicating copy number status of SHH subgroup known somatic driver genes CCND2, GLI2, MYCN, and PTEN.

We implemented molecular subtyping as follows:

1. MB tumors with methylation classification that contains “MB_SHH” are subtyped as SHH-activated medulloblastoma (MB, SHH)
2. MB tumors with “MB_G34_I”, “MB_G34_II”, “MB_G34_III”, and “MB_G334_IV” methylation classifications are subtyped as medulloblastoma group 3 (MB, Group3)
3. MB tumors with “MB_G34_V”, “MB_G34_VI”, “MB_G34_VII”, and “MB_G334_VIII” methylation classifications are subtyped as medulloblastoma group 4 (MB, Group4)
4. MB tumors with “MB_WNT” methylation classification are subtyped as WNT-activated MB (MB, WNT)
5. MB tumors with “MB_MYO” methylation classification are subtyped as medulloblastomas with myogenic differentiation (MB, MYO)

We classified MB, SHH subtype tumors using the following criteria:

1. *MB, SHH alpha*: sample has a high-confidence “MB_SHH_3” methylation classification, or patient had an age at diagnosis >= 2 years and harbored one of the following molecular alterations in tumor or germline:

- *MYCN*, *GLI2*, or *CCND2* amplification or sample TPM z-score >= 2 in tumor.
- A pathogenic or likely pathogenic germline variant in *ELP1* or *TP53*.
- A *TP53* hotspot mutation in tumor.
- Chromosome 9p gain or chromosome 17p loss in tumor.
2. *MB, SHH beta*: sample has a high-confidence “MB_SHH_1” methylation classification, or patient had an age at diagnosis < 5 years and harbored one of the following molecular alterations:

- A *KMT2D* loss of function variant.
- *PTEN* copy number loss or deep deletion, or sample TPM z-score < -2.
- Chromosome 2p or 2q gain.
3. *MB, SHH gamma*: sample has a high-confidence “MB_SHH_2” methylation classification, or patient had an age at diagnosis < 5 years and tumor harbored a chromosome 2p arm gain.
4. *MB, SHH delta*: sample has a high-confidence “MB_SHH_4” methylation classification, or patient had an age at diagnosis >= 10 years and harbored one of the following molecular alterations in tumor:

- a *DDX3X* or *SMO* loss-of-function mutation.
- a hotspot *TERT* or U1 snRNA gene mutation.
- Chromosome 14q arm loss.

##### Pineoblastomas

Pineoblastomas (PB) are classified as follows using high-confidence methylation classifications:

1. Pineoblastoma, MYC/FOXR2-activated (“PB_FOXR2” methylation classification)
2. Pineoblastoma, RB1-altered (“PB_RB1” methylation classification)
3. Pineoblastoma, group 1 (“PB_GRP1A” and “PB_GRP1B” methylation classifications)
4. Pineoblastoma, group 2 (“PB_GRP2” methylation classification)
5. All other pineoblastomas were classified as “PB, To be classified”

##### non-MB, non-ATRT Embryonal Tumors

Updates were made to non-MB, non-ATRT embryonal tumor subtyping as follows:

1. Embryonal tumors with multilayered rosettes and C19MC-altered (ETMR, C19MC-altered) were classified based on 1) high-confidence “ETMR_C19MC” methylation classification or 2) *TTYH1* gene fusion and either chromosome 19 amplification or *LIN28A* over-expression.
2. ETMR, not otherwise specified (NOS) were classified based on *LIN28A* over-expression and no *TTYH1* gene fusion.

### TP53 Alteration Annotation (tp53_nf1_score analysis module)

We classified TP53-altered high-grade glioma (HGG) samples as either *TP53* lost or *TP53* activated and incorporated these annotations into the molecular subtype framework. To support this classification, we used a previously published RNA-based *TP53* inactivation signature originally developed using TCGA pan-cancer cohorts [40]. We applied this to OpenPedCan RNA-seq data, stratified by library preparation type. This classifier was used in combination with genomic variant data, including consensus SNVs, CNVs, and structural variants (SVs), as well as curated reference databases cataloging somatic *TP53* hotspot mutations [41,42] and known functional domains [43] to annotate lost or activated status. Briefly, samples were annotated as *TP53* activated if they harbored either of two known gain-of-function mutations: p.R273C or p.R248W [44]. Samples were assigned *TP53* lost status under any of the following conditions: (i) presence of a hotspot *TP53* mutation listed in the IARC or MSKCC databases; (ii) detection of two distinct TP53 alterations (e.g., SNV, CNV, or SV) consistent with biallelic inactivation; (iii) presence of a single somatic *TP53* variant or a pathogenic germline variant associated with Li-Fraumeni syndrome (LFS) [45]; or (iv) presence of a germline *TP53* variant linked to LFS alongside a *TP53* inactivation classifier score >0.5 from matched RNA-seq data.

### Clinical data harmonization

To remain consistent with the Kids First data model and our previous OpenPBTA study [1], all clinical metadata was harmonized using the same data model. TARGET and TCGA metadata fields (e.g., `sample_type`, `composition`, `tumor_descriptor`, etc.) were harmonized to those of Kids First. Additional histology-related fields were created through OpenPedCan, following molecular subtyping: `integrated_diagnosis`, `harmonized_diagnosis`, and `cancer_group`. These fields were expanded from our previous study, to utilize the WHO 2021 CNS tumor classifications[36]. Any samples with molecular subtypes which did not match the initial `pathology_diagnosis` were reviewed with a board-certified molecular pathologist and updated accordingly.

### EFO, MONDO, and NCIT Mapping

We created a script to search ontology mappings by `cancer_group`. The `efo_code` represents the Experimental Factor Ontology (EFO) description available in European Bioinformatics Institute database, the `mondo_code` represents the Mondo Disease Ontology (MONDO) from an independent resource that aims to harmonize disease definitions, and the `ncit_code` represents the NCI Thesaurus (NCIt) reference terminology. Codes were automatically pulled based on text matching, manually reviewed, and can be found in **Supplemental Table 1**

#### Selection of independent samples (independent-samples analysis module)

For analyses that require all input biospecimens to be independent, we use the OpenPedCan-analysis <u>independent-samples</u> module to select only one biospecimen from each input participant. For each input participant of an analysis, the independent biospecimen is selected based on the analysis-specific filters and preferences for the biospecimen metadata, such as experimental strategy, cancer group, and tumor descriptor.

## Data Validation and Quality Control

All RNA-seq and WGS samples passed minimum quality thresholds, including ≥20 million total reads and ≥50% alignment for RNA-Seq, and ≥20X mean coverage for DNA sequencing **Supplemental Table 4**. Sample identity was confirmed using NGSCheckMate [46] and Somalier relate [47] to detect and exclude mismatched or contaminated samples.

We expanded upon the molecular subtyping modules from OpenPBTA to recover hallmark genomic and transcriptomic features known in pediatric tumors. These include: *KIAA1549::BRAF* fusions in low-grade gliomas, H3 K28M/I mutations in diffuse midline gliomas, H3 G35R/V mutations in diffuse hemispheric gliomas, somatic *TP53* mutations in high-grade gliomas, and MYCN amplification in neuroblastoma, for example.

All subtyping modules are version-controlled, containerized, and publicly available, and have undergone internal code review and validation by independent analysts. Where molecular features conflicted with original pathology labels, cases were reviewed with board-certified molecular pathologists, and integrated diagnoses were updated accordingly. This collaborative re-review process led to improved sample annotation and is fully documented in the molecular-subtype-pathology module.

To assess concordance between data types, we compared RNA-based and methylation-based molecular subtypes in medulloblastoma. As shown in [Table 1], we observed nearly 100% concordance, validating both experimental modalities and classifier accuracy. Notably, methylation classification identified one rare case (MB, MYO) not captured by the transcriptome-based MedulloClassifier.

**Table 1:** Medulloblastoma subtype concordance across experimental strategies. Comparison of medulloblastoma subtypes using methylation or RNA-Seq classification.

| Methylation Subtype | Group3 (RNA-Seq) | Group4 (RNA-Seq) | SHH (RNA-Seq) | WNT (RNA-Seq) |
| --- | --- | --- | --- | --- |
| MB_G34_II | 8 | 0 | 0 | 0 |
| MB_G34_III | 17 | 0 | 0 | 0 |
| MB_G34_IV | 8 | 0 | 0 | 0 |
| MB_G34_V | 0 | 6 | 0 | 0 |
| MB_G34_VI | 0 | 4 | 0 | 0 |
| MB_G34_VII | 0 | 30 | 0 | 0 |
| MB_G34_VIII | 0 | 34 | 0 | 0 |
| MB_MYO | 1 | 0 | 0 | 0 |
| MB_SHH_1 | 0 | 0 | 11 | 0 |
| MB_SHH_2 | 0 | 0 | 4 | 0 |
| MB_SHH_3 | 0 | 0 | 2 | 0 |
| MB_SHH_4 | 0 | 0 | 7 | 0 |
| MB_WNT | 0 | 0 | 0 | 18 |

**Table 2:**
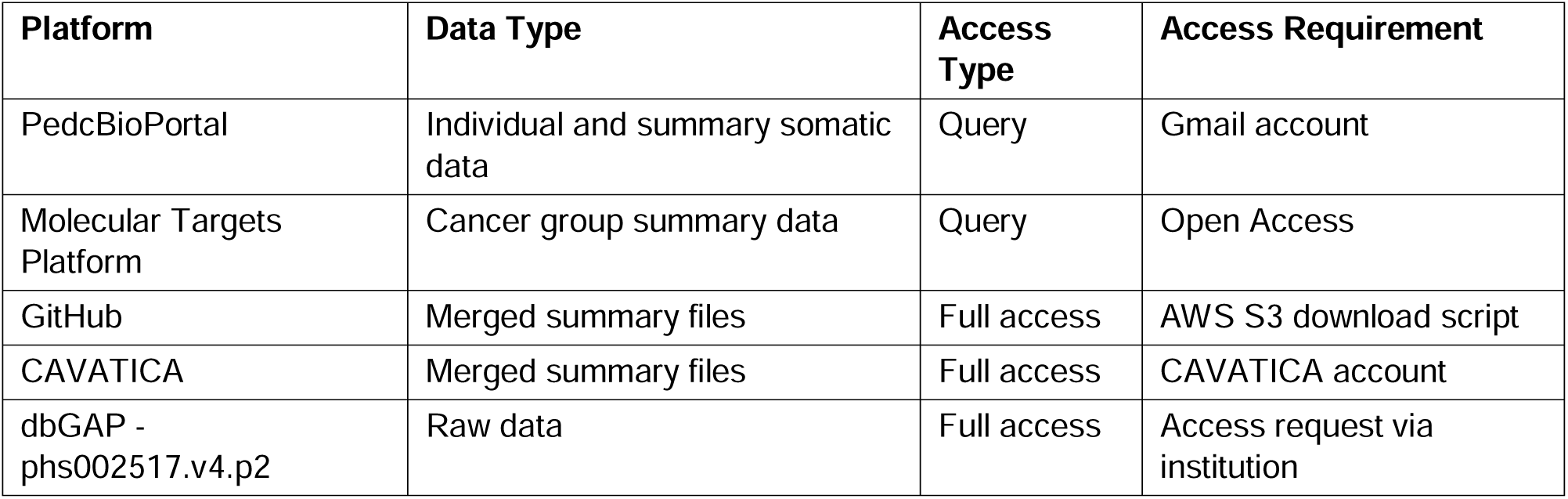
OpenPedCan Data Availability. OpenPedCan data is available on multiple platforms with varying access requirements.

| Platform | Data Type | Access Type | Access Requirement |
| --- | --- | --- | --- |
| PedcBioPortal | Individual and summary somatic data | Query | Gmail account |
| Molecular Targets Platform | Cancer group summary data | Query | Open Access |
| GitHub | Merged summary files | Full access | AWS S3 download script |
| CAVATICA | Merged summary files | Full access | CAVATICA account |
| dbGAP - phs002517.v4.p2 | Raw data | Full access | Access request via institution |

In addition to verifying known findings, OpenPedCan modules support pediatric cancer discovery and translation. The reproducibility of these results is further supported by their reuse across studies, > 100 Zenodo downloads, GitHub forks, and independent analysis pipelines.

Together, these validation measures—spanning sample QC, molecular feature recovery, cross-platform concordance, and expert review—ensure that OpenPedCan is a robust, reproducible, and reusable resource for the pediatric cancer research community.

## Ethics and Consent Statement

This study did not generate new sequencing data. All previously-published raw data were obtained through Database of Genotypes and Phenotypes (dbGAP) access requests with patients consented as “General Research Use (GRU)” or “Disease-Specific (Pediatric Cancer Research)”. OpenPedCan integrates only summary-level outputs (e.g., gene expression matrices, mutation calls) that are designated for GRU by the data custodians. No protected health information (PHI), raw sequencing files, or individually identifiable clinical metadata are distributed as part of this project.

## Re-use potential

OpenPedCan represents a valuable resource, not only by significantly extending OpenPBTA to include more than 5,000 additional patients and 6,000 tumors, but also by adding a number of new “omic” data types not previously included, such as methylation arrays, miRNA-Seq, proteomics, and normal tissue RNA-Seq. OpenPedCan also serves as a community resource whose outputs and/or code can be leveraged directly to ask research questions or serve as an orthogonal validation dataset. By providing this data in a harmonized manner, we enable investigators to reduce the financial and time-related costs associated with their analyses, which would otherwise total years of project hours and over $50,000 in data analysis alone [48]. We encourage re-use of the data, ideas and suggestions for improving the data or adding analyses, and/or direct code contributions through a pull-request.

## Availability of source code and requirements

Project name: The Open Pediatric Cancer (OpenPedCan) Project

Project home page: https://github.com/d3b-center/OpenPedCan-analysis

Archived Source code: https://zenodo.org/records/15750097

Operating system(s): Platform independent

Programming languages: R, Python, bash

Other requirements: CAVATICA is required to run all primary Kids First workflows. All downstream OpenPedCan workflows can be run using the Docker image at pgc-images.sbgenomics.com/d3b-bixu/openpedcanverse:latest. Most workflows run efficiently on local or cloud machines with 16–64 GB RAM. The most memory-intensive module runs on a 64 GB instance at <$2 per run.

License: CC-BY 4.0

Primary analyses were performed using Gabriella Miller Kids First pipelines and are listed in the methods section. Analysis modules were either initially developed within https://github.com/AlexsLemonade/OpenPBTA-analysis [1], were modified, and/or created anew within the https://github.com/d3b-center/OpenPedCan-analysis publicly available repository.

Software versions are documented in **Supplemental Table 5**.

## Data Availability

### Datasets

The datasets supporting this study are available as follows: The TARGET dataset is available in dbGAP under phs000218.v23.p8 [49]. The GMKF Neuroblastoma dataset is available in dbGAP under phs001436.v1.p1[50]. The Pediatric Brain Tumor Atlas data (PBTA), containing the subcohorts OpenPBTA, Kids First PBTA (X01), Chordoma Foundation, MI-ONCOSEQ Study, PNOC, and DGD is available in dbGAP under phs002517.v4.p2 [51] or in the Kids First Portal (kidsfirstdrc.org). The raw Genotype-Tissue Expression (GTEx) dataset is available in dbGAP under phs000424.v9.p2 and publicly available at https://gtexportal.org/home/. The Cancer Genome Atlas (TCGA) dataset is available in dbGAP under phs000178.v11.p8 [52].

Merged summary files for the latest release of OpenPedCan are openly accessible in <u>CAVATICA</u> or via `download-data.sh` script in the https://github.com/d3b-center/OpenPedCan-analysis repository. Cancer group summary data from release v11 are visible within the NCI’s pediatric <u>Molecular Targets Platform</u>. Cohort, cancer group, and individual data are visible within <u>PedcBioPortal</u>. An overview of the OpenPedCan data availability is summarized in [**Table 22**].

## Acknowledgments

We are incredibly grateful to each patient and family for donating tissue and associated metadata and clinical data to their respective consortia. This project has been funded in whole or in part with Federal funds from the National Cancer Institute, National Institutes of Health, under Contract No. 75N91019D00024, Task Order No. 75N91020F00003 (DMT, JLR, SJD, JMM, ST, AF, ACR). The content of this publication does not necessarily reflect the views or policies of the Department of Health and Human Services, nor does mention of trade names, commercial products or organizations imply endorsement by the U.S. Government. The authors also wish to thank the anonymous private investors to the Children’s National Hospital Brain Tumor Institute who have supported this work. We thank Rocky Breslow for GitHub actions contributions and Rust Turakulov for contributing to methylation data analysis.

## Author Contributions

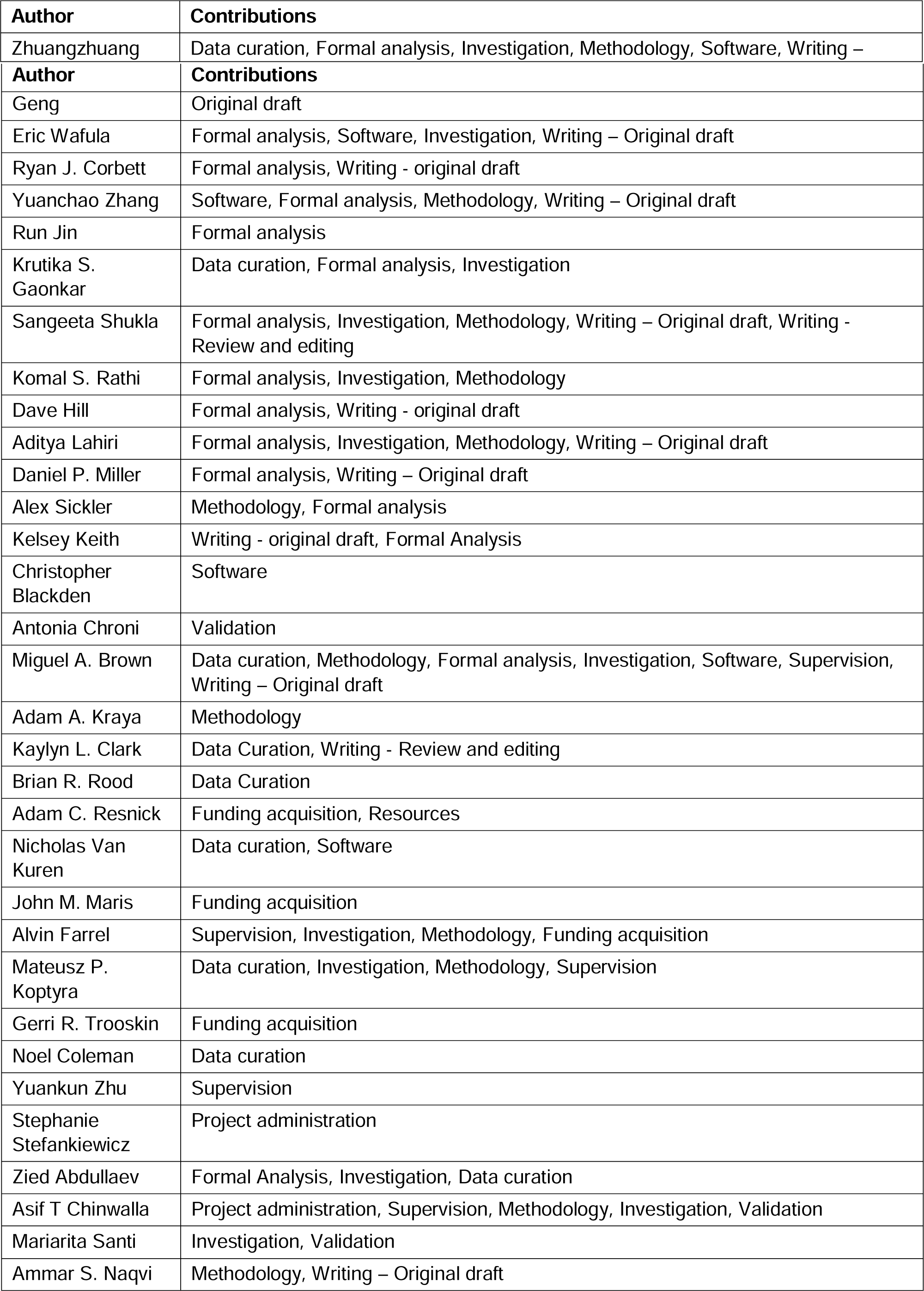
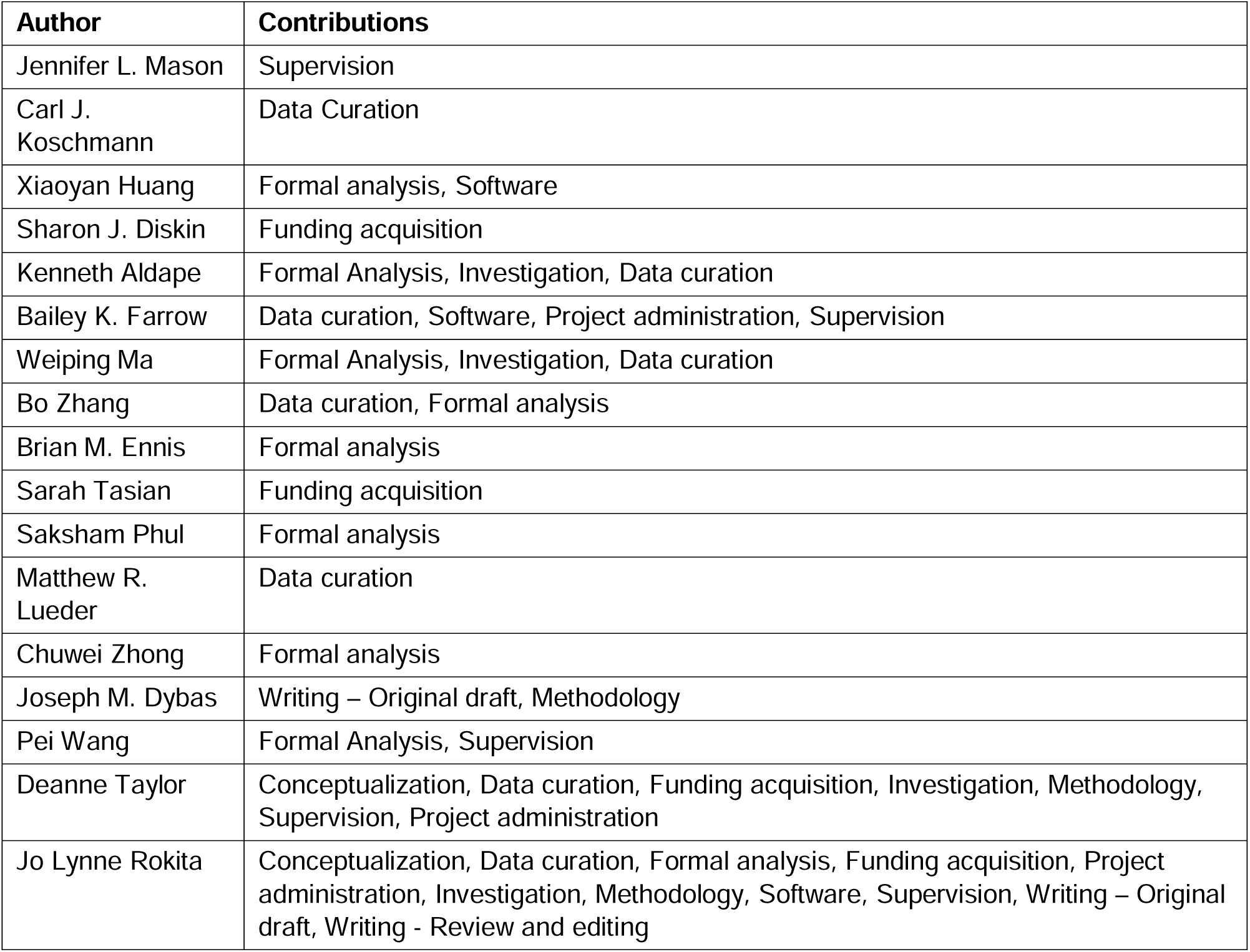

